# Independent evolution of lateral inhibition mechanisms in different lineages of land plants: MpFEW RHIZOIDS1 miRNA-mediated lateral inhibition controls rhizoid cell patterning in *Marchantia polymorpha*

**DOI:** 10.1101/854869

**Authors:** Anna Thamm, Timothy E Saunders, Liam Dolan

## Abstract

Lateral inhibition patterns differentiated cells during development in bacteria, metazoans and land plants. Tip-growing rhizoid cells develop among flat epidermal cells in the epidermis of the early diverging land plant *Marchantia polymorpha*. We show that the majority of rhizoid cells develop individually but some develop in linear, one-dimensional clusters of between two and seven rhizoid cells in wild type plants. The distribution of rhizoid cells can be accounted for within a simple model of lateral inhibition. The model also predicted that, in the absence of lateral inhibition, rhizoid cell clusters would be two-dimensional with larger clusters than those formed with lateral inhibition. Rhizoid differentiation in *Marchantia polymorpha* is positively regulated by the ROOT HAIR DEFECTIVE SIX-LIKE1 (MpRSL1) basic Helix Loop Helix (bHLH) transcription factor which is directly repressed by the FEW RHIZOIDS1 (MpFRH1) miRNA. To test if MpFRH1 miRNA acts during lateral inhibition we generated loss-of-function mutants that did not produce the MpFRH1 miRNA. Two-dimensional clusters of rhizoids develop in Mp*frh1^loss-of-function (lof)^* mutants as predicted by the model for plants that lack lateral inhibition. Furthermore, clusters of up to nine rhizoid cells developed in the Mp*frh1^lof^* mutants compared to a maximum number of seven observed in wild type. The higher steady state levels of MpRSL1 mRNA in Mp*frh1^lof^* mutants indicate that MpFRH1-mediated lateral inhibition involves the repression of Mp*RSL1* activity. Together the modelling and genetic data indicate that the pattern of cell differentiation in the *M. polymorpha* epidermis is consistent with a lateral inhibition process in which MpFRH1 miRNA represses MpRSL1. This discovery suggests that novel mechanisms of lateral inhibition may operate in different lineages of land plants, unlike metazoans where the conserved Delta-Notch signaling system controls lateral inhibition in diverse metazoan lineages.

## INTRODUCTION

Spatial arrays of differentiated cells form during the development of multicellular organisms. The patterning of different cell types during animal development involves lateral inhibition, a process in which an individual cell instructs adjacent cells to acquire an identity that is different from the fate of the instructing cell. In metazoans, lateral inhibition involves cell-to-cell signalling carried out by the Delta-Notch ligand-receptor pair (reviewed in Artavanis-Tsakonas et al., 1999; Bray, 2006; Perrimon et al., 2012). Neither Delta nor Notch proteins are present in plants and other mechanisms of lateral inhibition operate in the angiosperm *Arabidopsis thaliana* (The Arabidopsis Genome Initiative, 2000; Wigge and Weigel, 2001; Bowman et al., 2017). Computational modelling suggests that the production of EPIDERMAL PATTERNING FACTOR2 (EPF2) peptide by meristemoids inhibits meristemoid development in adjacent cells during guard cell development in leaves (Horst et al., 2015; Lee et al., 2015; Simmons and Bergmann, 2016). The cell-to-cell movement of Myb transcriptional repressor proteins pattern trichomes (leaf hair cells) among pavement epidermal cells in the *A. thaliana* shoot epidermis (Schellmann et al., 2002). A similar mechanism is required for the maintenance of patterning in the root epidermis (Lee and Schiefelbein, 2002; Savage et al., 2008). The demonstration that there are at least two mechanisms of lateral inhibition in *A. thaliana* alone suggests that, unlike metazoans, diverse mechanisms of lateral inhibition likely function among land plants.

Rhizoid cells develop from fields of equivalent cells in the epidermis of the liverwort *Marchantia polymorpha,* an early diverging land plant. Flat epidermal cells occupy the space between rhizoid cells in wild type plants. The RSL1 basic-Helix-Loop-Helix transcription factor is necessary and sufficient for rhizoid cell development (Menand et al., 2007; Catarino et al., 2016; Proust et al., 2016). Rhizoid cells do not develop in plants harbouring loss-of-function Mp*rsl1* mutations and flat epidermal cells develop in their place (Proust et al., 2016). Conversely plants harbouring Mp*RSL1* gain-of-function mutations develop supernumerary ectopic rhizoids in place of flat epidermal cells (Proust et al., 2016). MpRSL1 promotes the expression of Mp*FRH1* which encodes a 21-nucleotide microRNA (Honkanen et al., 2018). Evidence that MpFRH1 represses rhizoid cell development by negatively regulating MpRSL1 activity include (i) FRH1 miRNA directly targets RSL1 mRNA for DICERLIKE1-mediated cleavage and (ii) the rhizoidless phenotype of Mp*FRH1* overexpression is suppressed by co-expressing an MpFRH1 miRNA-resistant version of MpRSL1 (Honkanen et al., 2018). Therefore, MpRSL1 and MpFRH1 act together in a network with feedback where the transcription factor is an activator of rhizoid cell development and the miRNA is a repressor of rhizoid cell development.

Since MpRSL1 and MpFRH1 constitute an activator-repressor network with negative feedback, we hypothesized that the MpFRH1 miRNA could be involved in lateral inhibition during the patterning of rhizoid cells in the epidermis *M. polymorpha* (Honkanen et al., 2018). To test this hypothesis, we first modelled the effect of lateral inhibition on the pattern of rhizoid cell spacing in wild type conditions. This revealed that the spatial distribution of rhizoid cells in wild type was consistent with lateral inhibition being involved in patterning. We then ran the model in the absence of lateral inhibition to predict the rhizoid cell distribution that would be observed in a plant in which lateral inhibition did not operate. To test if MpFRH1 miRNA was involved in lateral inhibition we characterised rhizoid cell distribution in Mp*frh1^loss-of-function (lof)^* mutants. The distribution of rhizoid cells in Mp*frh1* mutants was consistent with the predicted distribution from the model without lateral inhibition. These data indicate that MpFRH1 miRNA acts in the lateral inhibition of rhizoid cell development during epidermal development in *M. polymorpha*.

## RESULTS

### Linear clusters of rhizoid cells develop in wild type *Marchantia polymorpha* gemma epidermis

The *M. polymorpha* gemma epidermis comprises flat epidermal cells that surround rhizoid cells that produce a tip growing projection that is involved in rooting function. To determine if lateral inhibition is involved in the patterning of rhizoid cells, we characterised their spatial arrangement on the dorsal surface of wild type gemmae (Tak-1 and Tak-2). Rhizoid cells develop individually and in clusters in wild type gemmae (Figure 1; Table 1). The majority (∼70%) of rhizoid cells develop individually, surrounded by flat epidermal cells in wild type. In the Tak-2 background, approximately 30% of rhizoids develop in clusters of between 2 and 6 rhizoid cells (Table 1); 18.8% 7.3%, 2.6%, 1.0%and 1.0% of rhizoids formed in clusters of 2, 3, 4, 5 and 6 rhizoid cells, respectively. Most rhizoid cells in clusters are arranged in a one-dimensional, linear array, like beads on a string. Rhizoid cells at either end contact one neighbouring rhizoid cell and internal rhizoid cells always contact two rhizoid cells. Consequently, in a linear cluster of three rhizoid cells, there are two rhizoid cells on either end of the cluster with a single rhiziod cell between each of these end initials. Therefore, such a cluster of three develops an average of 1.33 rhizoid cell neighbours ((1 + 2 + 1)/3 = 1.33). The rhizoid cells in 24 of the 25 3-rhizoid cell clusters were one-dimensional. The rhizoid cells in a single 3-rhizoid cell cluster contacted each other and had an average of 2 rhizoid cell neighbours (two-dimensional). The average number of rhizoid cell neighbours in 3-rhizoid cell clusters was therefore 1.36, (SD = 0.13; n= 25). The mean number of rhizoid cell neighbours increases as cluster size increases. For example in a one-dimensional cluster of 4-rhizoid cells there is an average of 1.5 rhizoid cell neighbours ((1+2+2+1)/4 = 1.5). Rhizoid cells in 12 of 13 four-cell cluster were arranged in one-dimensional clusters while rhizoid cells were arranged in a two-dimensional cluster in a single case. Therefore the mean rhizoid cell neighbour number in four-cell cluster is 1.57 (SD = 0.27; n= 13). We observed four 5-rhizoid cell clusters and three 6-cell cluster, and rhizoid cells were arranged in a one-dimension in all seven. Therefore the average rhizoid cell neighbour number was = 1.6 in 5-cell cluster (SD = 0; n= 4 clusters) and 1.66 in 6-cell cluster (SD = 0; n=3) (Figure 1). There was a single, two-dimensional, 7-cell cluster in the Tak-1 background (Figure S1). Hence, for such one-dimensional rhizoid arrays with *n* cells, the average rhizoid cell neighbour number = 2-2/n. The SD is close to zero indicating that this pattern is consistent in each cluster class. In total, 93.5 % of rhizoid clusters in wiltype are arranged in a one-dimension and only 6.5 % are arranged in a two dimensions (n= 46, Figure 4c). Based on those findings, the maximum number of rhizoid neighbour cells is generally two and clusters comprise one-dimensional linear arrays of rhizoid cells. These data show that wild type rhizoid cells form either individually or in linear clusters where each rhizoid cell has either one or two neighbours and not more than two. This is consistent with the hypothesis that lateral inhibition operates in the patterning of rhizoid cells in the epidermis.

**Figure 1.**
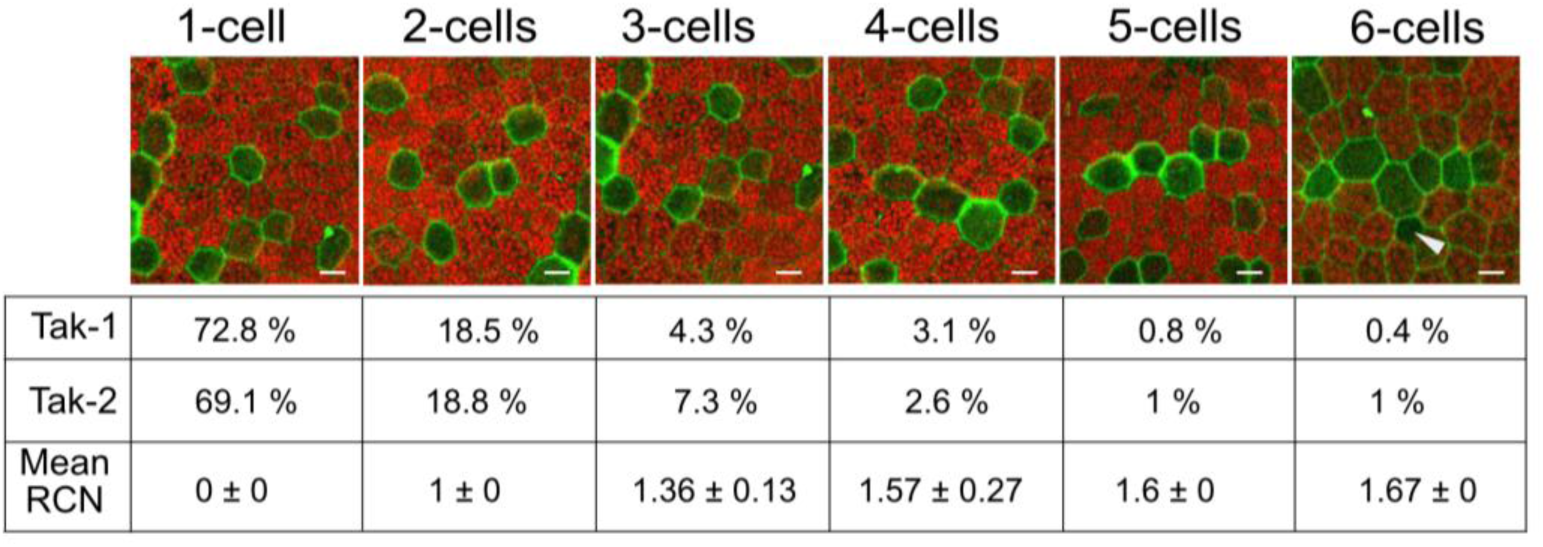
Organisation of rhizoid clusters in the *M. polymorpha* gemma epidermis. Representative examples of individual rhizoid cells and clusters with between 2 and 6 rhizoid cells on mature gemmae of Tak-1 and Tak-2 wild type accessions. Chlorophyll autofluorescence is shown in red, cell walls of rhizoid cells are stained with propidium iodide (green). Tabular presentation of the frequencies of each size class of rhizoid cells cluster. The values represent the frequency of rhizoid cluster developing of each class as a percentage of the total number of rhizoid cell cluster. The mean rhizoid cell neighbour number (RCN) for each size class is represented with the standard deviation. (n [Tak-1] = 22 gemmae, n [Tak-2] = 23). Scale bar 20 µm, arrowhead indicates the position of an oil body cell which is smaller than a rhizoid cell.

**Table 1.** Frequency of individual rhizoid cells and rhizoid cluster in wildtype, Mp*frh1^lof^* and Mp*RSL1^GOF^* mutants.

| genotype | 1 | 2 | 3 | 4 | 5 | 6 | 7 | 8 | 9 | ≥10 | n cluster | n gemmae |
| --- | --- | --- | --- | --- | --- | --- | --- | --- | --- | --- | --- | --- |
| Tak-1 | 72.5% | 18.4% | 4.3% | 3.1% | 0.8% | 0.4% | 0.4% |  |  |  | 255 | 22 |
| Tak-2 | 69.1% | 18.8% | 7.3% | 2.6% | 1.0% | 1.0% |  |  |  |  | 191 | 23 |
| *Mpfrh1*<sup>lof-7</sup> | 47.1% | 23.2% | 12.3% | 9.0% | 4.5% | 1.3% |  | 0.6% | 1.9% |  | 155 | 42 |
| *Mpfrh1*<sup>lof-11</sup> | 61.4% | 23.8% | 7.4% | 4.8% | 1.6% | 0.5% | 0.5% |  |  |  | 189 | 27 |
| *Mpfrh1*<sup>lof-27</sup> | 63.2% | 23.0% | 9.8% | 0.6% | 1.1% | 1.1% | 0.6% | 0.6% |  |  | 174 | 22 |
| *Mpfrh1*<sup>lof-34</sup> | 48.0% | 40.0% | 8.0% | 4.0% |  |  |  |  |  |  | 75 | 9 |
| *Mpfrh1*<sup>lof-36</sup> | 69.4% | 19.4% | 6.9% | 1.4% | 2.8% |  |  |  |  |  | 72 | 8 |
| *MpRSL1*<sup>GOF</sup> | 58.6% | 14.1% | 13.1% | 5.1% | 3.0% | 1.0% | 2.0% |  |  | 3.0% | 99 | 10 |

### Modelling predicts that lateral inhibition in involved in the patterning of rhizoid cells in wild type gemmae

A model for the development of rhizoid cells in the gemma epidermis was developed. We initiated a lattice of cells with seeds for rhizoid cells at similar density to measured experimentally. The model had two rules. Rule (1): For each rhizoid, at each simulation step a probability P determined whether the cluster stops expanding (and stays fixed at that cell number for remainder of simulation) or (1-P) that a new neighbouring cell becomes a rhizoid cell. P was estimated from the data from wild type clusters. To incorporate new rhizoid cells in wild type conditions we applied the Rule (2): the new rhizoid cell had to be a neighbour to precisely one existing rhizoid cell (no fewer and no more) (Figure 2a). No other rules or parameters were incorporated into the model. We analysed the number and distribution of clusters produced by the model. With this simple rule, the model replicated the observed distributions of rhizoid cell cluster sizes and the rhizoid cell neighbour frequency as observed experimentally (Figure 2a, b). All rhizoid cell clusters that developed were linear – each rhizoid cell in the cluster developed either one rhizoid cell neighbour (if located at the end of a cluster) or two rhizoid cell neighbours (if located away from cluster ends) and the SD for mean rhizoid cell neighbour number was 0 in each size class. Thus, simple lateral inhibition is consistent with our observed patterns of rhizoid cell development and also with the invariant neighbour number.

**Figure 2.**
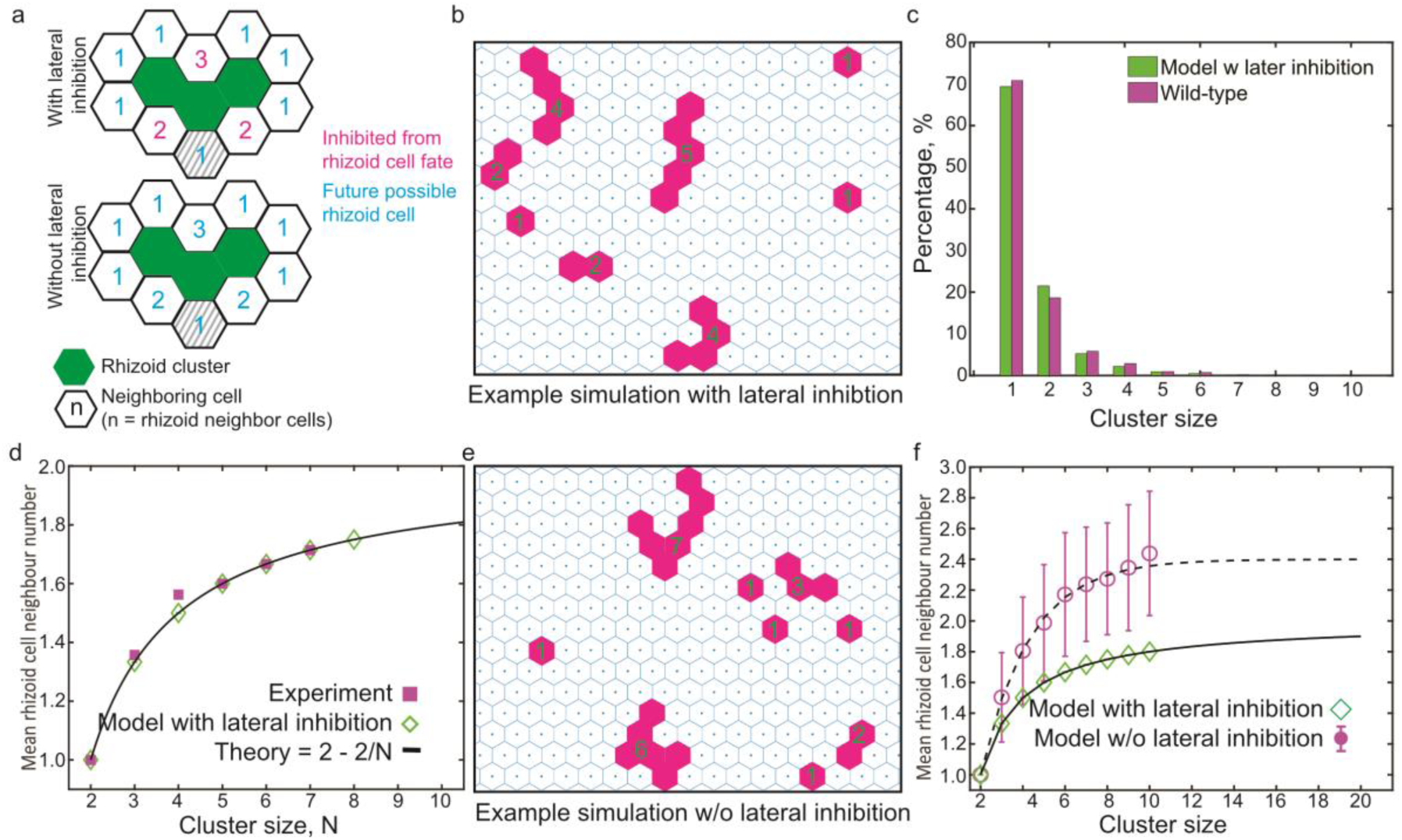
Modeling indicates that lateral inhibition can account for the patterning of rhizoid cells in the *M. polymorpha* gemma. a. Schematic of lateral inhibition model of rhizoid development. (Top) Lateral inhibition restricts the addition of a new cell to a rhizoid cluster to those only neighbouring a single rhizoid cell. (Bottom) In the absence of lateral inhibition, any cell can differentiate as a rhizoid cell. Grey cell represents a new possible rhizoid cell that is not adjacent to the end of the rhizoid cluster. We exclude such a possibility in the simulations, but including this does not alter the statistical results (Supplementary Information). b. Output of the lateral inhibition model showing rhizoid cell distribution (magenta) and flat epidermal cells (white) in the gemma epidermis. In the presence of lateral inhibition, each cluster forms a linear array with cells arranged like beads on a string. c. Validation of lateral inhibition model by comparing the frequency distribution of rhizoid cell clusters produced by the model (green, n = 500 simulations) and those observed in wild type gemma epidermis (magenta, “experiment”, n = 45 gemmae). d. Validation of lateral inhibition model by comparing the relationship between mean rhizoid cell neighbour number and rhizoid cluster size as predicted by the model (green) with data from wild type gemmae (magenta). e. Output of the model without lateral inhibition, showing the distribution of rhizoid cells (magenta) and flat epidermal cells (white) in the gemma epidermis. In the absence of lateral inhibition, clusters can develop in two-dimensional arrays. f. Predicted relationship between mean rhizoid cell neighbour number and rhizoid cluster size in the presence and absence of lateral inhibition. The lower curve, with green points, indicates the output from the model that includes lateral inhibition. The upper curve with magenta points, indicates the output of the model without lateral inhibition. The main differences are than the mean rhizoid cell number is greater in the absence of lateral inhibition than with lateral inhibition for e each rhizoid cell cluster size and while these values are invariant in the presence of lateral inhibition they are variable in the absence of lateral inhibition.

We next considered the scenario whereby rule 2 was relaxed so that a new cell can develop next to one *or more* existing rhizoid cells; *i.e.* we remove the lateral inhibition requirement. The resulting pattern did not resemble the pattern of rhizoid cells in wild type plants (Figure 2c); clusters were two-dimensional instead of one-dimensional. In the absence of lateral inhibition, larger rhizoid cell clusters developed than in wild type due to merging of neighbouring clusters. Furthermore, the variability in the number of rhizoid cell neighbours, measured by SD, for rhizoid cell cluster larger than 3 was always greater than 0, the value predicted for the simulation with lateral inhibition. Other more complex models can be considered. However, our simple Boolean-like lateral inhibition model is consistent with the hypothesis that lateral inhibition functions during the patterning of rhizoids; a rhizoid cell can develop next to flat epidermal cells or a single rhizoid but never more than one rhizoid cell.

With our given *P*, clusters of 6 or more cells are not seen in the wild type simulations where lateral inhibition is active, which is consistent with what is observed in wild type plants. However, in our simulations with the relaxed Rule 2, we noticed that occasionally clusters can merge; in the absence of lateral inhibition, two rhizoid cell clusters are not inhibited from adding a joining cell. In this case, larger clusters form, comprising up to 12 cells in the model, but these are rare (observed once in over 1000 simulations). The model predicts both an increase in the rhizoid cell neighbour number and an increase in the standard deviation in cluster neighbours. Furthermore, these clusters have rounder shapes than the elongated, linear wild type structures (Figure 2e). These arrangements are only rarely observed in the models in which lateral inhibition operates in wild type plants. These modelling results indicate that if lateral inhibition controls the spacing of rhizoid cells in *M. polymorpha*, loss of this mechanism will cause development of two-dimensional rhizoid cell clusters and rare large clusters. We emphasise that these quantitative model predictions require no extra parameters; they are simply dependent on the presence/absence of lateral inhibition.

### Two dimensional clusters developed in Mp*frh1* mutants

If MpFRH1 miRNA mediates lateral inhibition during rhizoid cell development, the model predicted that two-dimensional rhizoid clusters would develop in Mp*frh1^lof^* mutants while one-dimensional rhizoid clusters would develop in wild type. To determine if MpFRH1 miRNA is involved in lateral inhibition we generated mutants in which the entire Mp*FRH1* miRNA coding sequence was deleted. We used a pair of single guide RNAs (sgRNAs) 569 bp apart to delete the MpFRH1 gene using CRISPR/Cas9 technology (Sugano et al., 2014). sgRNA1 was 20 bp long and was complementary to a sequence between 404bp – 384 bp 5’ of the Mp*FRH1* miRNA sequence (sgRNA1) and sgRNA2 was 20 bp long and complementary to a sequence between 164 bp – 184 bp 3’ of the Mp*FRH1* miRNA sequence (Figure 3a). Five independent mutant lines were identified in which the entire Mp*FRH1* miRNA sequence was deleted. There were deletions of 267 bp, 587 bp, 589 bp and 573 bp in the Mp*FRH1* gene of Mp*frh1-*7, Mp*frh1*-27, Mp*frh1*-34 and Mp*frh1*-36 mutants respectively. In Mp*frh1*-11, a 780 bp deletion removed the Mp*FRH1* sequence and a 202 bp sequence inserted 5’ of the Mp*FRH1* sequence. Mp*FRH1* miRNA was not detected (Figure 3b) in any of the five deletion mutants consistent with the hypothesis that each harboured complete loss-of-function Mp*frh1* mutations. Consequently, the mutants were designated Mp*frh1^lof-7,^* Mp*frh1^lof-11^,* Mp*frh1^lof-27^,* Mp*frh1^lof-34^* and Mp*frh1^lof-36^* (where lof is loss-of-function and distinguishes them from Mp*FRH1^GOF^* lines which harbour gain-of-function mutations).

**Figure 3.**
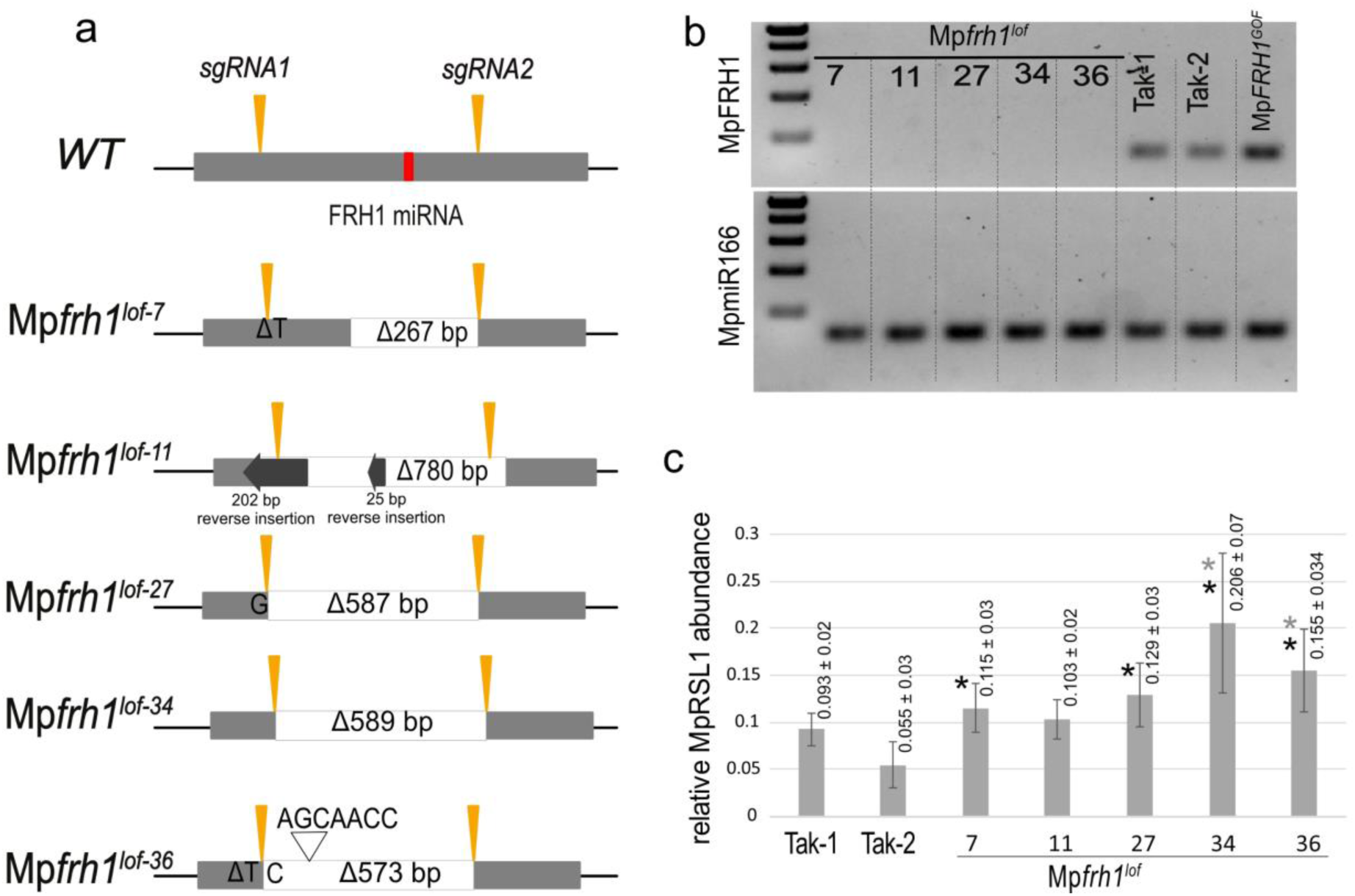
Genomic organisation of Mp*frh1* mutants and expression of Mp*RSL1* and Mp*FRH1* in Mp*frh1* loss of function mutants. a. Genomic organisation of the Mp*FRH1* gene in wild type (WT) and Mp*frh1^lof^* mutants. The sequence for the 1.2kb MpFRH1 transcript is represented with a grey box. The 21 nucleotides corresponding to the MpFRH1 miRNA are indicated in red. The orange arrow heads indicate the position of the two single guide RNAs (sgRNAs) used to generate deletion mutants. sgRNA1 is complementary to a sequence that is between 384 and 404 bp 5’ of the MpFRH1 sequence. SgRNA2 is complementary to a sequence that is 164-184 3’ of the MpFRH1 sequence. The entire Mp*FRH1* miRNA sequence was deleted in each of the mutants. In Mp*frh1^lof-7^* there was a 267bp deletion between sgRNA1 and sgRNA2 and T deletion. In Mp*frh1^lof-11^* there was a deletion of 780bp and reverse insertions of 202bp and 26bp. In Mp*frh1 ^lof-27^* there was a 587bp deletion and a G insertion. In Mp*frh1^lof-34^* there was a 589bp deletion. In Mp*frh1^lof-36^* there was a 573bp deletion and substitution of T into C at the sgRNA1 target site and a 7bp insertion. b. Amplification of MpFRH1 miRNA (upper row) and MpmiR166 control (lower row) using stem loop PCR on RNA isolated from 7-day old thalli grown from gemmae in Mp*frh1^lof^* mutants, Tak-1, Tak-2, and Mp*FRH1^GOF^* mutant. c. Amplification of MpRSL1 mRNA from RNA isolated from in 7-day old thalli grown from gemmae in wild type (Tak-1, Tak-2) plants and Mp*frh1^lof^* mutants; mRNA levels were normalized to the housekeeping gene Mp*APT1*. Three biological replicates per genotype, except for Mp*frh1^lof-7^*(n=2). There were at least three technical replicates per biological replicate. Grey asterisk: significant difference to Tak-1, black asterisk: significant difference to Tak-2. P<0.05

To determine if the pattern of rhizoids in Mp*frh1^lof^* mutants was as predicted by the model in the absence of lateral inhibition, the number of rhizoid cell neighbours was measured in rhizoid clusters in each of the mutants. The model predicted that the number of rhiziod cell neighbours in clusters would be greater than in wild type. As predicted by the model, the mean rhizoid cell neighbour number is greater in the mutants than in wild type rhizoid cell clusters for each cluster size class (Figure 4b, Figure 5). In all five Mp*frh1^lof^* mutants, the number of neighbours in clusters of three rhizoid cells was greater than a 1.36 (the average number of neighbours in a cluster of three rhizoid cells in wild type). Furthemore while the SD was close to 0 in wild type indicating that there was little variation in the mean number of neighbour cells per cluster, the SD was always greater than 0 in the mutants. For example, there were on average 1.4 (SD = 0.21) rhizoid cell neighbours in clusters of three rhizoid cells in Mp*frh1^lof-7^*, 1.45 (SD = 0.26) rhizoid cell neighbours in Mp*frh1^lof-27^*, 1.48 (SD = 0.28) rhizoid cell neighbours in Mp*frh1^lof-11^,* 1.56 (SD=0.34) in Mp*frh1^lof-34^* and 1.6 (SD=0.37) in Mp*frh1^lof-^*^36^ (Figure 5a, e). The same trend held for clusters of four rhizoid cells where the number of rhizoid cell neighbours was 1.57 (SD = 0.27) in wild type (1.63 for Tak-1 (SD = 0.35), 1.5 for Tak-2 (SD = 0)). Clusters containing four rhizoid cells developed an average 1.73 (SD = 0.37) rhizoid cell neighbours in Mp*frh1^lof-7^*, 1.61 (SD =0.33) in Mp*frh1^lof-11^*, 1.67 (SD = 0.29) in Mp*frh1^lof-34^* mutants (Figure 5b, e,). Clusters of five rhizoid cells in Mp*frh1^lof-27^* developed 2.2 (SD = 0.28) rhizoid cell neighbours compared to 1.6 (SD = 0) in wild type (Figure 5c, e). Consequently rhizoid cell clusters are two dimensionsal in Mp*frh1^lof^* mutants rather than one-dimensional as in in wild type. Remarkably, our simple model predicted similar shifts in both the mean and standard deviation under loss of the purported lateral inhibtion (Figure 2).

**Figure 4.**
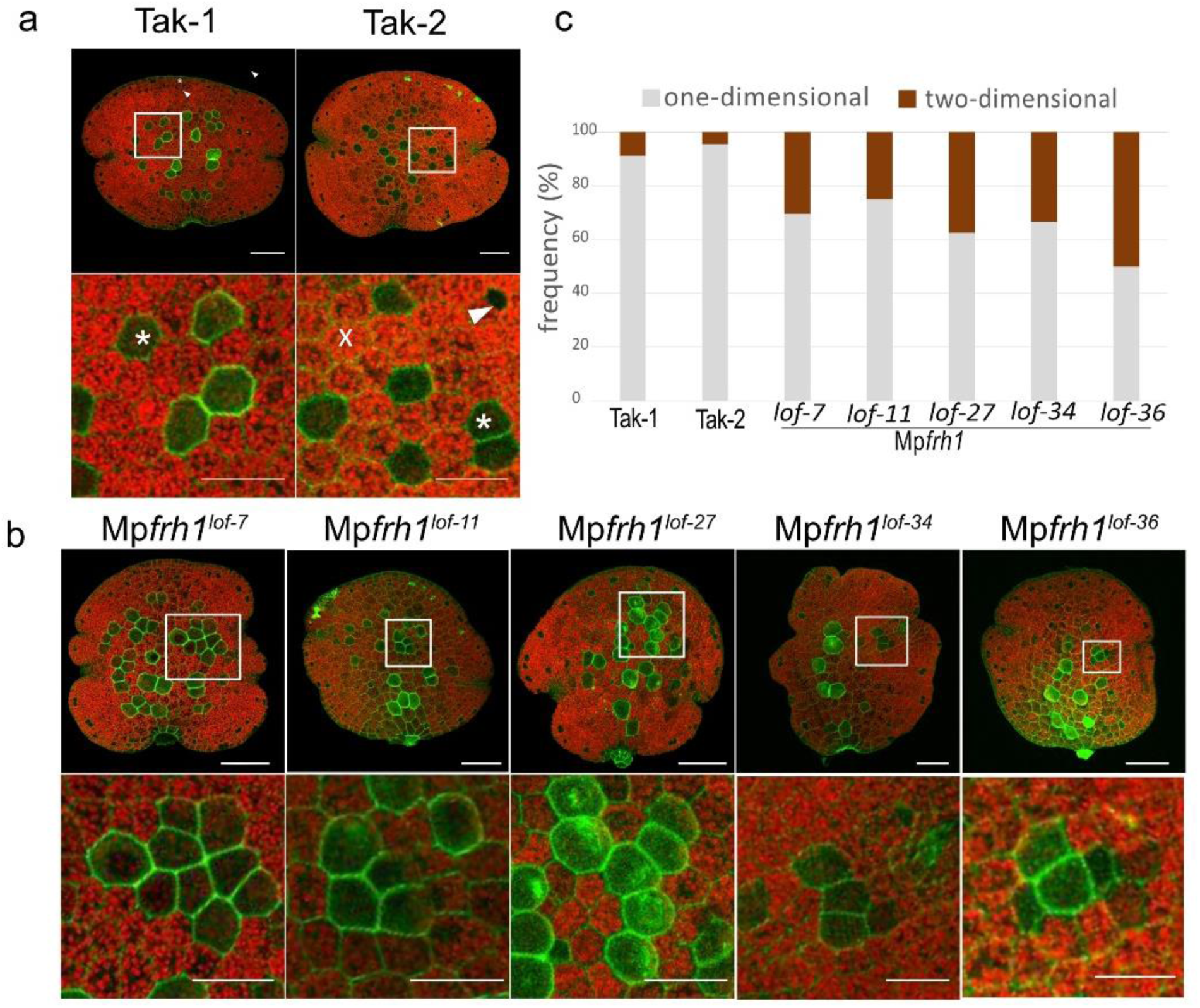
Two-dimensional rhizoid cell clusters develop in Mp*frh1^lof^* and Mp*RSL1^GOF^* mutants. a. Rhizoid patterning in gemmae of wild type (Tak-1, Tak-2). Upper row, overview of representative gemma (scalebar = 100 µm). Lower row shows the higher magnification of inset highlighted above, showing representative rhizoid clusters (Scalebar = 50µm). b. Rhizoid patterning in gemmae of Mp*frh1^lof-7^*, Mp*frh1^lof-11^*, Mp*frh1^lof-27^*, Mp*frh1^lof-34^*, Mp*frh1^lof-36^* mutants. Upper row, overview of representative gemma (scalebar = 100 µm). Lower row, magnification of highlighted square showing representative rhizoid cluster (scalebar = 50µm). c. Frequency (in %) of one-dimensional and two-dimensional rhizoid cluster in wildtype (Tak-1, Tak-2) and Mp*frh1^lof^* mutants. All rhizoid cluster larger than three cells were combined (n[Tak-1] = 23, n[Tak-2=23], n[Mp*frh1^lof-7^*] = 46, n[Mp*frh1^lof-11^*] = 28, n[Mp*frh1^lof-27^*] = 24, n[Mp*frh1^lof-34^*] = 9, n[Mp*frh1^lof-36^*]=8).

**Figure 5.**
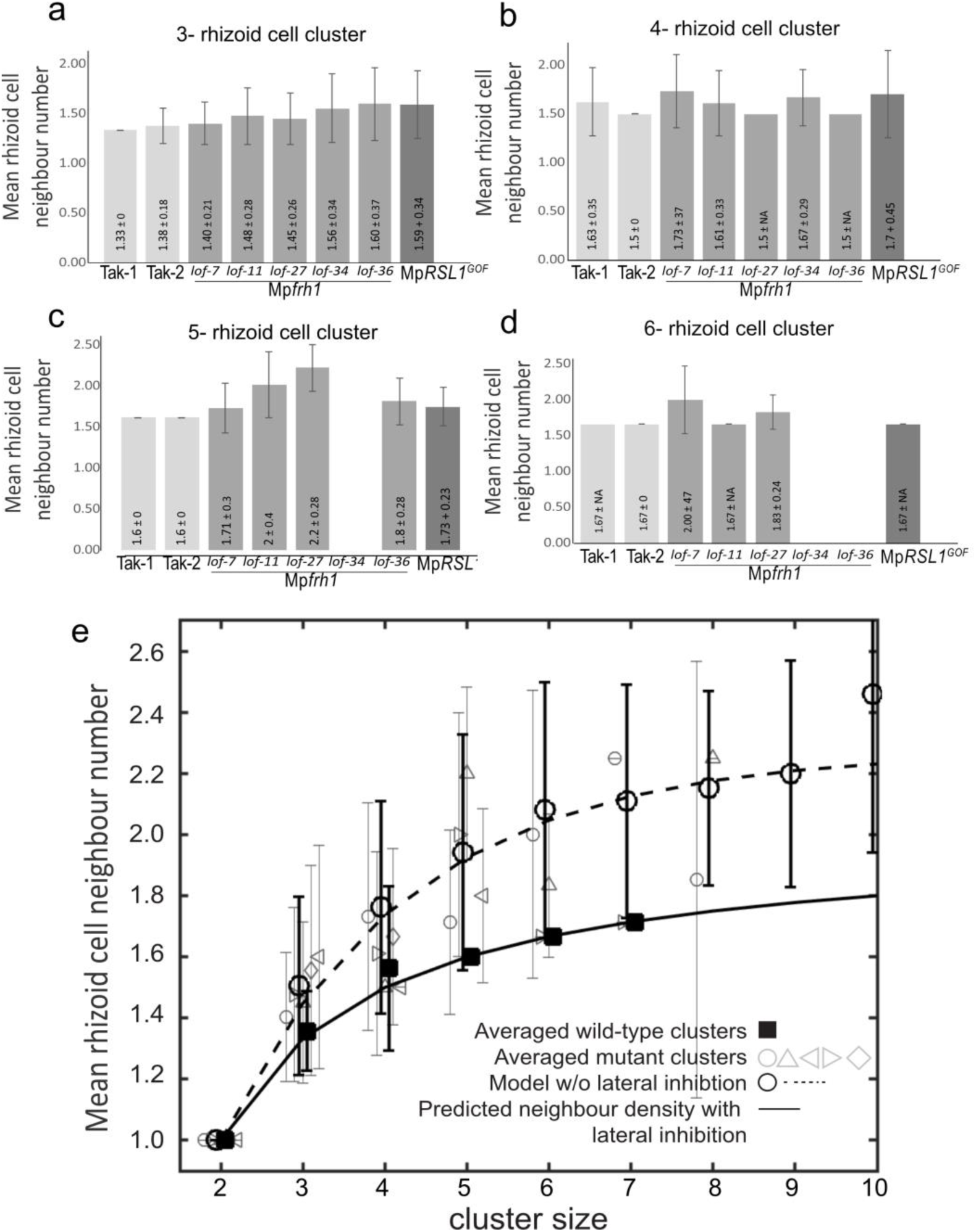
More rhizoid cell neighbours develop in Mp*frh1^lof^* and Mp*RSL1^GOF^* mutants. a-d. Mean rhizoid cell neighbour number in 3-rhizoid cell clusters (a), 4-rhizoid cell clusters (b), 5-rhizoid cell clusters (c), and 6-rhizoid cell clusters (d) in wild type (Tak-1, Tak-2), Mp*frh1^lof-7^*, Mp*frh1^lof-11^*, Mp*frh1^lof-27^*, Mp*frh1^lof-34^*, Mp*frh1^lof-36^* and Mp*RSL1^GOF^*. e. The relationship between mean rhizoid cell neighbour numbers and cluster size predicted by the model with lateral inhibition (solid line). The relationship predicted by the model without lateral inhibition (open circles, n= 500 simulations; dashed line represents fit to 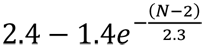 where *N* is cluster size). The experimentally observed relationship between mean rhizoid cell neighbour numbers and cluster size in wild type (filled black square) and Mp*frh1^lof^* mutants (Mp*frh1^lof-7^* = grey circle, Mp*frh1^lof-11^* = grey triangle pointing to right, Mp*frh1^lof-27^* = grey triangle pointing up, Mp*frh1^lof-34^* = grey diamond, Mp*frh1^lof-36^* = grey triangle pointing to left). Errorbars are SD.

**Figure 6.**
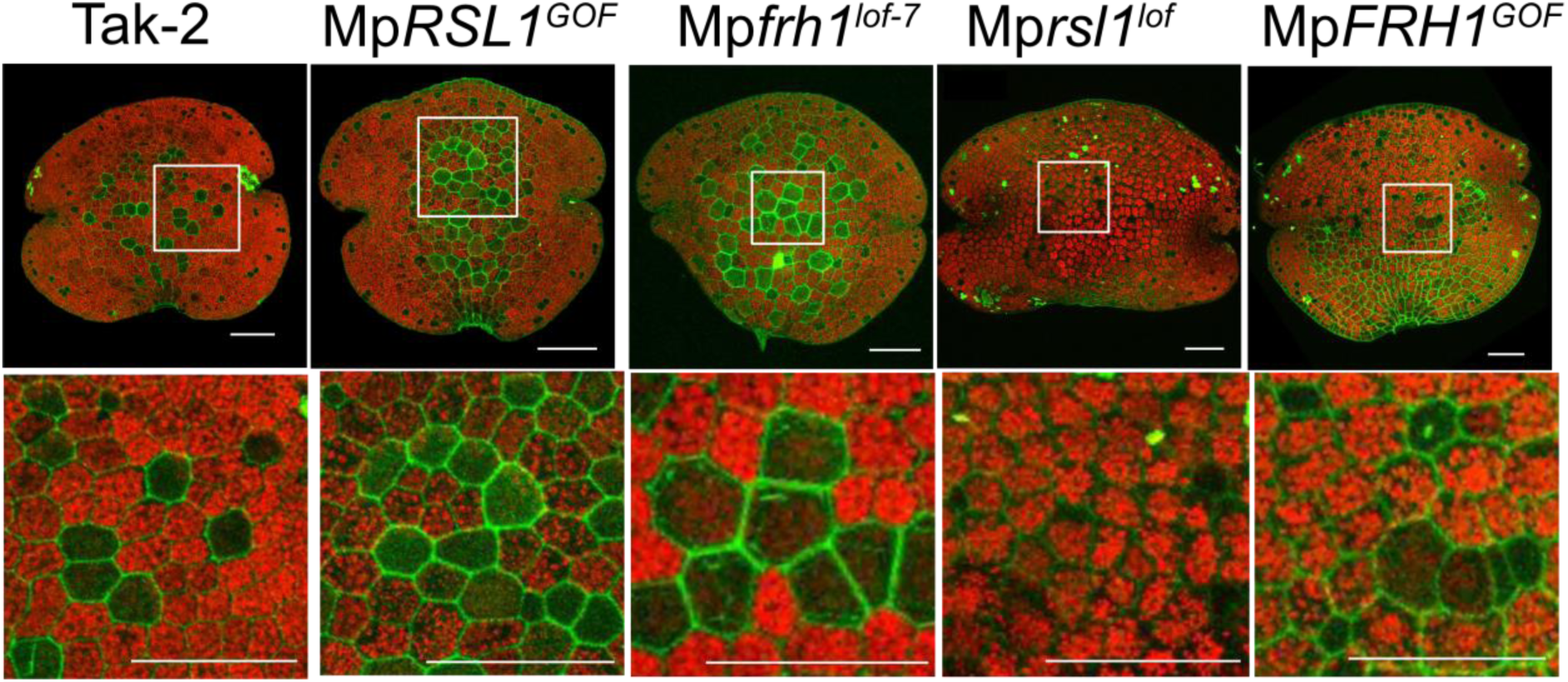
Larger rhizoid cell clusters with altered shape develop in Mp*frh1^lof^* and Mp*RSL1^GOF^* mutants than in wild type. Upper row shows distribution of rhizoid cells and rhizoid cell clusters on representative mature gemmae of wild type (Tak-2), Mp*RSL1^GOF^*, Mp*frh1^lof^*^-7^, Mp*rsl1^lof^*, Mp*FRH1^GOF^*mutants. Scalebar = 100 µm. The lower row shows the higher magnification of the inset highlighted on the image in the row above. There are 12 rhizoid cells in the Mp*RSL1^GOF^* rhizoid cell cluster and 8 cells in the Mp*frh1^lof^*^-7^ rhizoid cell cluster. There are no rhizoids in either the Mp*rsl1^lof^* or Mp*FRH1^GOF^* mutants. Chlororphyll autofluorescence is red, and cell walls of rhizoids are stained with propidium iodide (green). Scalebar = 50 µm. White arrowhead indicates the location of oil body cells.

To confirm that the increased number of rhizoid cell neigbours resulted in an increased frequency of two-dimensional clusters, as predicted by the model, we measured the frequencecy of one-dimensional and two-dimensional rhizoid clusters in Mp*frh1^lof^* mutants. 92-96% of rhizoid cluster are one-dimensional and only 4-8% of rhizoid cluster are two-dimensional in wild type. By contrast, 32% of rhizoid cluster are two-dimensional in Mp*frh1^lof^* mutants. This includes 30% of rhizoid clusters in Mp*frh1^lof-^* 7 (14 of 46), 25% in Mp*frh1^lof-11^* (7 of 21), 37% in Mp*frh1^lof-27^* (9 of 24), 33% in Mp*frh1^lof-34^* (3 of 9) and 50% in Mp*frh1^lof-36^* (4 of 8 cluster)(Figure 4c). The higher frequency of two-dimensional clusters in Mp*frh1^lof^* than in wildtype is consistent with the hypothesis that Mp*FRH1* activity is required for lateral inhibition during rhizoid development. The phenotypes of larger rhizoid cell neighbour number and higher frequency of two-dimensional rhizoid cell clusters in Mp*frh1^lof^* mutants than in wild type are consistent with the hypothesis that Mp*FRH1* is active in lateral inhibition during rhizoid cell development.

If MpFRH1 miRNA acts as an inhibitor during lateral inhibition, the model predicted that rhizoid clusters with up to 9 rhizoid cells should develop infrequently in Mp*frh1^lof^* mutants. Such clusters form from the merger of adjacent clusters; a process which is inhibited in wild type conditions. The maximum number of rhizoid cells in clusters was 7 in wild type (Table 1, Figure 2c). Rhizoid cell clusters with more than 7 cells were found in Mp*frh1^lof^* mutants (Table 1, Figure 4b). A single 8-rhizoid cell cluster and three 9-rhizoid cell clusters were observed in Mp*frh1^lof-7^* and a single 8-rhiziod cell cluster was found in Mp*frh1^lof-27.^* The observation that rare clusters containing up to 9 rhizoid cells developed in Mp*frh1^lof^* mutants is consistent with the hypothesis that MpFRH1 acts as an inhibitor in lateral inhibition during epidermal develoment.

### MpFRH1-mediated lateral inhibition acts by repressing Mp*RSL1* activity

Since MpFRH1 miRNA targets the MpRSL1 mRNA, we hypothesized that the loss of MpFRH1 function would result in an increase in the steady state levels of Mp*RSL1* mRNA. mRNA was isolated from thallus that developed from whole gemmae grown for 7 days. Mp*RSL1* mRNA abundance was normalised to the Mp*APT1* housekeeping gene (Saint-Marcoux et al., 2015). MpRSL1 mRNA is more abundant in Mp*frh1^lof^* mutants than in wild type. Steady state levels of MpRSL1 mRNA were 2-2.5 times more abundant in Mp*frh1^lof-36^* and Mp*frh1^lof-34^* than in wild type (Figure 3c). Steady state levels of MpRSL1 mRNA were approximately 1.5 times higher in Mp*frh1^lof-7^,* Mp*frh1^lof-11^* and Mp*frh1^lof-27^* mutants than in wild type. In an independent experiment with fewer replicates, the steady state levels of MpRSL1 mRNA were 3.7, 2.7, 4.23-, 3.77-, and 1.6-fold-higher in Mp*frh1^lof-7^,* Mp*frh1^lof-11^*, Mp*frh1^lof-27^* Mp*frh1^lof-34^* and Mp*frh1^lof-36^*, respectively, than in wild type (see Supplementary Figure2). These higher steady state levels of MpRSL1 mRNA in Mp*frh1^LOF^* mutants than wild type is consistent with the hypothesis that MpFRH1 targets MpRSL1 mRNA during lateral inhibition.

If MpFRH1-mediated lateral inhibition operates by repressing MpRSL1, overexpression of MpRSL1 should overcome the inhibitory effect of MpFRH1. If true, ectopic overexpression of MpRSL1 would increase the mean number of rhizod cell neighbours in clusters and that larger clusters of rhizoid cells would develop than in wild type. That is, the rhizoid cluster phenotypes of the Mp*RSL1^GOF^* and Mp*frh1^lof^* mutants would be similar. 3-rhizoid cell clusters in Mp*RSL1^GOF^* mutants developed an average of 1.59 (SD = 0.34, n = 13) neighbouring rhizoid cells, compared to 1.36 neighbours in wild type (SD = 0.13) (Figure 5a). 4-rhizoid cell clusters in Mp*RSL1^GOF^* mutants developed 1.7 (SD = 0.45, n = 5) rhizoid cell neighbours compared to 1.58 in wild type (SD =0.28) (Figure 5b). 5-rhizoid cell clusters in Mp*RSL1^GOF^* mutants developed 1.73 (SD = 0.23) neighbours, compared to 1.6 in wild type (SD = 0) (Figure 4c). The larger number and variability of rhizoid cell neighbours in Mp*RSL1^GOF^* mutants than in wild type indicate that ectopic overexpression of Mp*RSL1* causes similar phenotypic defecst in rhizoid clusters as in Mp*frh1^lof^* mutants. These data are consistent with the hypothesis that MpFRH1 miRNA represses Mp*RSL1* activity during lateral inhibition as rhizoids develop.

The model indicates that the maximum cluster size would be greater than 6 in the absence of lateral inhibition. If lateral inhibition were inactive in Mp*RSL1^GOF^* mutants, the model predicts that there would be more than 6 rhizoid cells in some clusters that develop in plants that ectopically overexress Mp*RSL1*. We counted the number of rhizoid cells in clusters in Mp*RSL1^GOF^* mutants. While the maximum number of rhizoid cells in wild type clusters was 7, there were up to 21 rhizoid cells in a single cluster in Mp*RSL1^GOF^* plants (Figure 1b, Supplement Figure 3). We observed three clusters greater than ten cells, included a single 10-cell cluster, a 12-cell cluster and a 21-cell cluster (summarzised to >10 cell cluster in Table1). This confirmes that repression of Mp*RSL1* occurs during lateral inhibition. Taken together, the data reported here demonstrates that MpFRH1 is a repressor that acts during lateral inhibition and MpFRH1 miRNA-mediated lateral inhibition acts by repressing Mp*RSL1* during the development of rhizoid cells in the *M. polymorpha* epidermis.

## DISCUSSION

We report that MpFRH1 miRNA represses rhizoid cell development through lateral inhibition in the *M. polymorpha* epidermis. Rhizoid cells develop individually or as one-dimensional clusters of between 2 and 7 rhizoid cells in the wild type epidermis. There is a high frequency of single rhizoid cells and progressively fewer clusters of 2, 3, 4, 5, 6 and-7 rhizoid cells. The spatial arrangement and distribution of rhizoid cells can be accounted for by a simple model that includes lateral inhibition. This model allowed us to predict the phenotype of rhizoid cell patterns that would develop in a mutant in which lateral inhibition was absent. It predicted that rhizoid cells would develop in two-dimensional clusters with larger clusters possible compared to wild type where clusters would be one-dimensional with a maximum of six cells in plants that lacked lateral inhibition. We showed that the rhizoid cell phenotypes that develop in plants that lack the MpFRH1 miRNA are those predicted by the model for an epidermis developing without lateral inhibition; clusters were two-dimensional and there were nine cells in the largest rhizoid cell cluster. These combined modelling and genetic data indicate that lateral inhibition mediated by the MpFRH1 miRNA is required for the patterning of rhizoid cells in *M. polymorpha*.

We propose that the development of rhizoid cells involves two distinct phases. The first phase involves the specification of individual rhizoid cells from a field of equivalent cells. The probability of a rhizoid cell being specified could be determined by stochastic variation in the expression levels of MpRSL1; higher levels would favour rhizoid cell development and relatively lower levels would favour flat epidermal cell development.

MpRSL1 then induces expression of the MpFRH1 miRNA in the developing rhizoid cell. MpFRH1 activity represses rhizoid cell development in adjacent cells through lateral inhibition and these cells develop as flat epidermal cells. Repeated rounds of rhizoid initiated during a competence period results in the development of rare adjacent rhizoid cells which form linear clusters, and the shape of these clusters is determined by lateral inhibition.

Our modelling and experimental data suggest that being adjacent to two rhizoid cells represses rhizoid cell development. By contrast, being adjacent to a single rhizoid cell is not sufficient to repress rhizoid development. This suggests that there is a threshold of repressor activity. Being adjacent to one rhizoid cell exposes an adjacent cell to sub-threshold levels of repressor while being adjacent to two rhizoid cells exposes a cell to repressor levels above this threshold and represses differentiation. A repressive signal from a rhizoid cell to adjacent cells represses rhizoid cell development if it reaches a threshold level in the receiving cell. MpFRH1 activity is required for this repressive signal. There are at least three potential modes of signalling. First, the repressive signal could be MpFRH1 miRNA itself. miRNAs can be mobile and it has been suggested that movement of miRNAs can account for the establishment of tissue boundaries in organs (Carlsbecker et al., 2010). Second, the repressive signal could be produced as a result of MpFRH1 miRNA activity. Since MpRSL1 mRNA is the only confirmed target of MpFRH1, the production of this hypothetical signal would likely also be MpRSL1-dependent (Honkanen et al., 2018). Third, the repressive signal could be transmitted by cell contact through an unknown mechanism. The amount of contact could be proportional to total shared cell surface area shared between neighbouring cells. Accordingly, sharing one cell face would expose the cells to sub-threshold levels of repressor while sharing a cell face with two rhizoid cells would expose the cell to repressor levels above this threshold. Since MpRSL1 mRNA is the only confirmed target of MpFRH1, the production of this hypothetical cell-contact signal would also be MpRSL1-dependent.

While the molecular mechanisms of lateral inhibition are different between plants (Myb repressor-, EDP2- and MpFRH1-dependent) and metazoans (Delta-Notch) there are similarities in the underlying logic of the process. For example, the Delta ligand represses neuroblast development during Notch-Delta signalling in Drosophila (reviewed in Bray, 2006). However, Delta is expressed on future neuroblasts and represses neuroblast development in adjacent cells by non-cell autonomous signalling mediated by the Notch receptor located on the neighbouring cell. That is, the repressor acts at a distance from its site of synthesis to repress the rhizoid cell identity in adjacent cells. This logic operates in Myb repressor- and EDP2-mediated lateral inhibition in *A. thaliana*. Similar logic operates in MpFRH1-mediated lateral inhibition in *M. polymorpha*. MpFRH1 is a repressor of rhizoid cell development that it is expressed in the developing rhizoid cells. Furthermore, our data indicate that MpFRH1 miRNA represses rhizoid cell identity in adjacent cells that would otherwise differentiate as flat epidermal cells in wild type. This leads to the hypothesis that MpFRH1 expression in developing rhizoid cells acts non-cell autonomously to repress rhizoid cell development in epidermal cells next to rhizoid cells. While the molecular mechanism of lateral inhibition is different between plants and animals, the regulatory similarities suggest that there may be underlying similarities in control logic.

The MpRSL1-MpFRH1 mechanism of lateral inhibition that controls the patterning of rhizoid cells is liverwort-specific. MpFRH1 is a liverwort-specific miRNA and the miRNA target sequence has been identified in the RSL genes of many liverwort taxa but has not been found in RSL genes from any other lineage of land plants (Honkanen et al., 2018). This indicates that the mechanism of lateral inhibition in the liverwort lineage is entirely different from the two mechanisms that has been described among the angiosperms (Myb repressor and EPF2). Since the MpFRH1-lateral inhibition mechanism is restricted to the liverworts it suggests that different lateral inhibition mechanisms control the patterning of tip growing rooting cells, rhizoids and root hairs in liverworts and angiosperms respectively. It also demonstrates that entirely different mechanisms can operate to control lateral inhibition in early diverging groups of land plants (hornworts and mosses) than in angiosperms. This contrasts with animals in which the Notch-Delta mechanism of lateral inhibition is conserved among most metazoan lineages. We propose that different mechanisms of lateral inhibition evolved many times during the course of land plant evolution while a single mechanism has been conserved among metazoans.

## Methods

### Plant material and growth conditions

M. polymorpha accessions Tagaragaike-1 (Tak-1, male) and Tagaragaike-2 (Tak-2, female)(Ishizaki et al., 2008) were used as wild type and grown as reported in (Honkanen et al., 2016). Mp*RSL1^GOF^* were generated by Proust et al., 2016. For RNA extraction plants were grown on medium covered with cellulosic cellophane membrane (AA Packaging Limited, Preston, UK) before harvesting to avoid agar contamination.

### CRISPR/Cas9 knock out

CRISPR/Cas9 mutations were generated following the protocol described in (Sugano et al., 2014). A modified version of pMpGE_En03 harbouring a second sgRNA scaffolds with a preceding BsmBI restriction site under the pU6 promoter was kindly provided by Holger Breuninger, named pHB453. Primer oAT110 and oAT111 were used to introduce sgRNA1 into the BbsI site, primer oAT112 and oAT113 were used to introduce sgRNA2 into BsmBI site. MpFRH1 allele was amplified using primer oAT135 and pAT136 and subsequentially sequenced by Source BioScience.

### qPCR of MpRSL1 and MpAPT

Total RNA was extracted from 7-day old gemmae using Direct-Zol RNA MINIprep kit (Zymo Research) following manufactures instructions. For each line three biological replicates were extracted unless stated otherwise. To remove DNA, 3ug total RNA were treated with TURBO DNA-free kit following manufacturers instruction. cDNA synthesis was performed according to the First Strand cDNA Synthesis protocol from NEB (#M0368) using ProtoSript II Reverse Transcriptase and Murine RNase Inhibitor. MpRSL1 and MpAPT cDNA was amplified using the primer oAT173/oAT174 and oAT175/176, respectively (as previously reported in Honkanen et al, 2018). qRT-PCT was performed in the Applied Biosystems 7300 Real-Time PCR System (Life Technologies) with SensiMix SYBR Hi-ROX Kit (Bioline) following manufactures instructions. Three technical replicates per biological replicate were performed. qPCR data was first analysed using LinRegPCR v2012.0(Ruijter et al., 2009). The average N_0_ value was calculated. Relative RSL1 mRNA abundance was calculated by normalising the N_0_ of each replicate against the N_0_ value of the reference gene, MpAPT1.

### Stem loop PCR for MpFRH1 and MpmiR166

To amplify MpFRH1 and MpmiR166, RNA was extracted from 19 day old gemmae using miRVana miRNA Isolation Kit following manufacturer’s instructions. 1ug RNA was treated with DNase according to TURBO DNA-free kit following manufacturers instruction. Stem-loop PCR was carried out as described in (Varkonyi-Gasic et al., 2007) using a MpFRH1 specific primer (oAT218) or a MpmiR166 specific primer (oAT219). The reverse transcribed and extended miRNAs were amplified using PCRBIO Ultra Polymerase with a miRNA specific forward primer (oAT221 for MpFRH1, oAT222 for MpmiR166) and a universal reverse primer (oAT220). Amplicons were visualised on a 3% agarose gel containing SYBR Safe.

### Imaging

Cell walls were stained using propidium iodide (PI). Gemmae were incubated in 5ug/ml PI solution for (5-)10 minutes. PI solution was removed and gemmae washed twice with MilliQ water. Gemmae were mounted on a slide with heated 0.2% agar solution. Once Agar solidified gemmae were imaged immediately. Images were acquired with a Leica SP5 confocal laser microscope and the Leica Application Suite (LAS) software using either a Leica HCX PL Fluotar 10x/0.30 or HC PL APO 20x/0.75 IMM CORR CS2 lense. PI and chlorophyll were excited at 543 nm and emission was collected between 561 and 640 nm for PI and between 680-700nm for chlorophyll.

For Scanning Electron Micropscopy (SEM), gemmae were fixed in 100% dry methanol for 30 minutes and then washed twiced with 100% dry ethanol. Gemmae were stored in 100% dry ethanol at 4 °C upon processing. The samples were dried at the critical point using a Tousimis Autosamdri-815, then mounted on aluminium stubs (Agar Scientific) using double-sided carbon adhesive discs (Agar Scientific), and coated with a mixture of gold and palladium using an SC7640 sputter coater (Quorum Technologies). The samples were imaged immediately with a JSM-5510 scanning electron microscope (JEOL) at an operating voltage of 15kV.

### Modelling

The model was implemented as follows. (1) A grid of ∼400 hexagonal cells was defined. A predefined density of cells were selected as rhizoid precursors. (2) A probability P defined the probability of a rhizoid cluster to include an additional neighbouring cell. Each rhizoid cluster was tested. If a cluster did not add a new cell (probability 1-P), then that cluster was considered to have finished expansion. (3) The simulation performed step (2) iteratively until all clusters finished expansion. In this simple scenario, the probability of a cluster having *n* cells = (1-P)^n-1^.

When a rhizoid cluster is selected for expansion, two possible rules were considered. (1) No lateral inhibition; in this case, any neighbouring cell to the cluster can become integrated within the cluster. (2) With lateral inhibition; only cells neighbouring exactly one member of the rhizoid cluster can become integrated within the cluster. We also considered the restriction that the new rhizoid cell has to be a neighbour of an end cell of the cluster. This corresponds to four possible configurations ((i) lateral inhibition with only end joining, (ii) lateral inhibition with new cells allowed so long as only one neighbour in the rhizoid cluster, (iii) no lateral inhibition with only end joining, (iv) no lateral inhibition with new cells allowed so long as only one neighbour in the rhizoid cluster). The results presented in the paper for wild type and mutants correspond to cases (i) and (iv) respectively. However, using rules (ii) and (iii) do not alter the general results, though clusters become more irregular as branching can occur.

When lateral inhibition was present, two nearby rhizoid clusters cannot merge; any cell between two clusters would have a minimum of two rhizoid neighbours. In the absence of lateral inhibition, such a cell can take on the rhizoid cell fate. Hence, two clusters can merge to form a larger cluster. In this way, although P is unchanged in all simulation, larger clusters are possible in the absence of lateral inhibition.

The model was encoded in Matlab. All data is produced from averaging the average cluster distributions for 1000 simulations. At each decision step, a uniform random number between 0 and 1 was drawn using the function *rand* and tested against the value *P*.

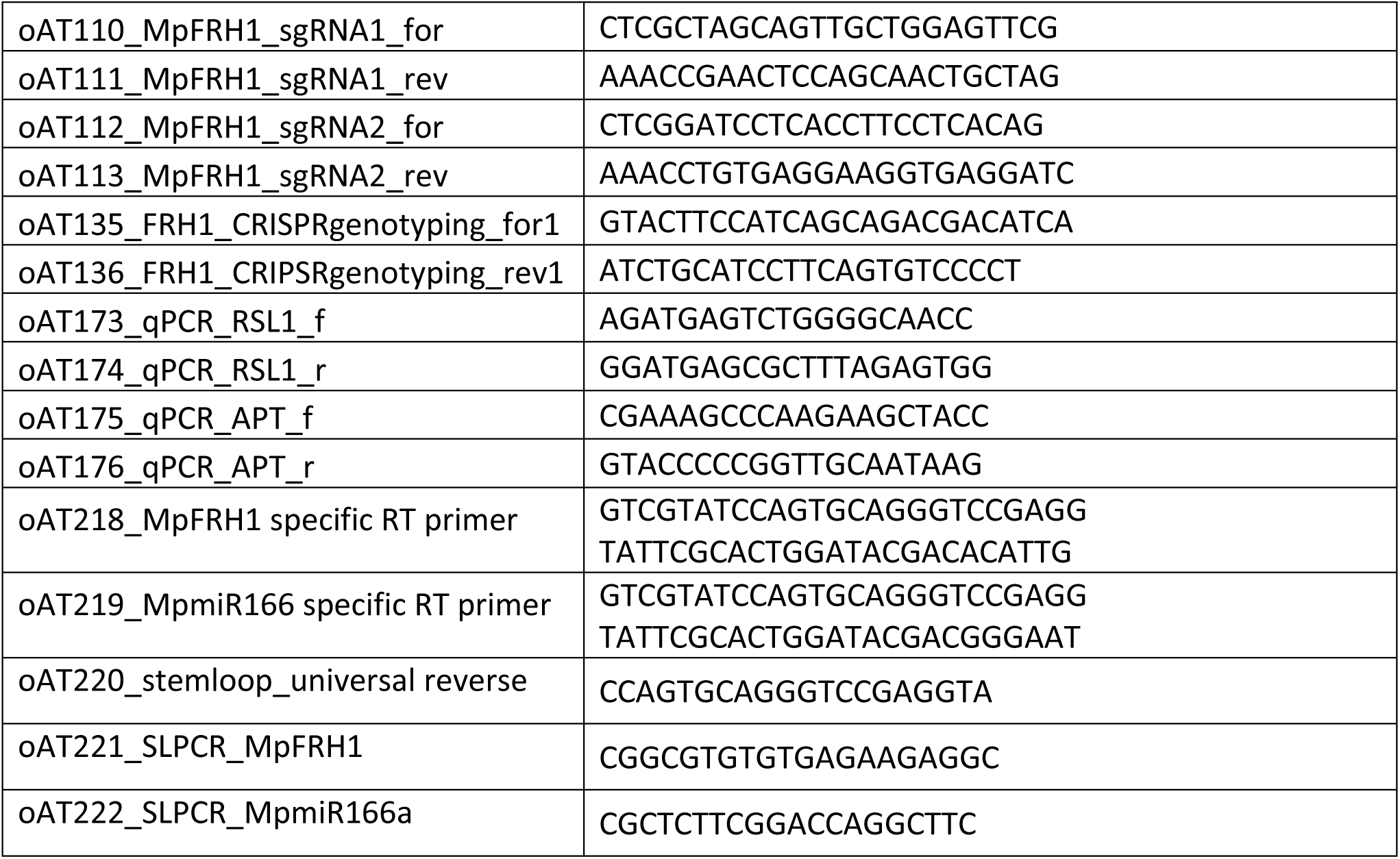
Primer list.

## Acknowledgment

Vector pHB453 was generated by Holger Breuninger after discussion with Julius Durr, Jose Gutierrez-Marcos, H. Puchta. T.E.S. and L.D. are grateful to the Kavli Institute, Santa Barbara, which supported their visits to the institute, supported in part by NSF Grant No. PHY-1748958, NIH Grant No. R25GM067110, and the Gordon and Betty Moore Foundation Grant No. 2919.01). T.E.S. acknowledges support from the Mechanobiology Institute and a Human Frontiers Young Investigator Grant. L.D. is funded by an European Research Council Advanced Grant (Denovo-P, contract 787613). A.T. was funded by Biotechnology and Biological Sciences Research Council (BBSRC) (BB/J014427/1) and Edward Penley Abraham Cephalosporin Scholarship.

---

**Supplementary Figure 1.**
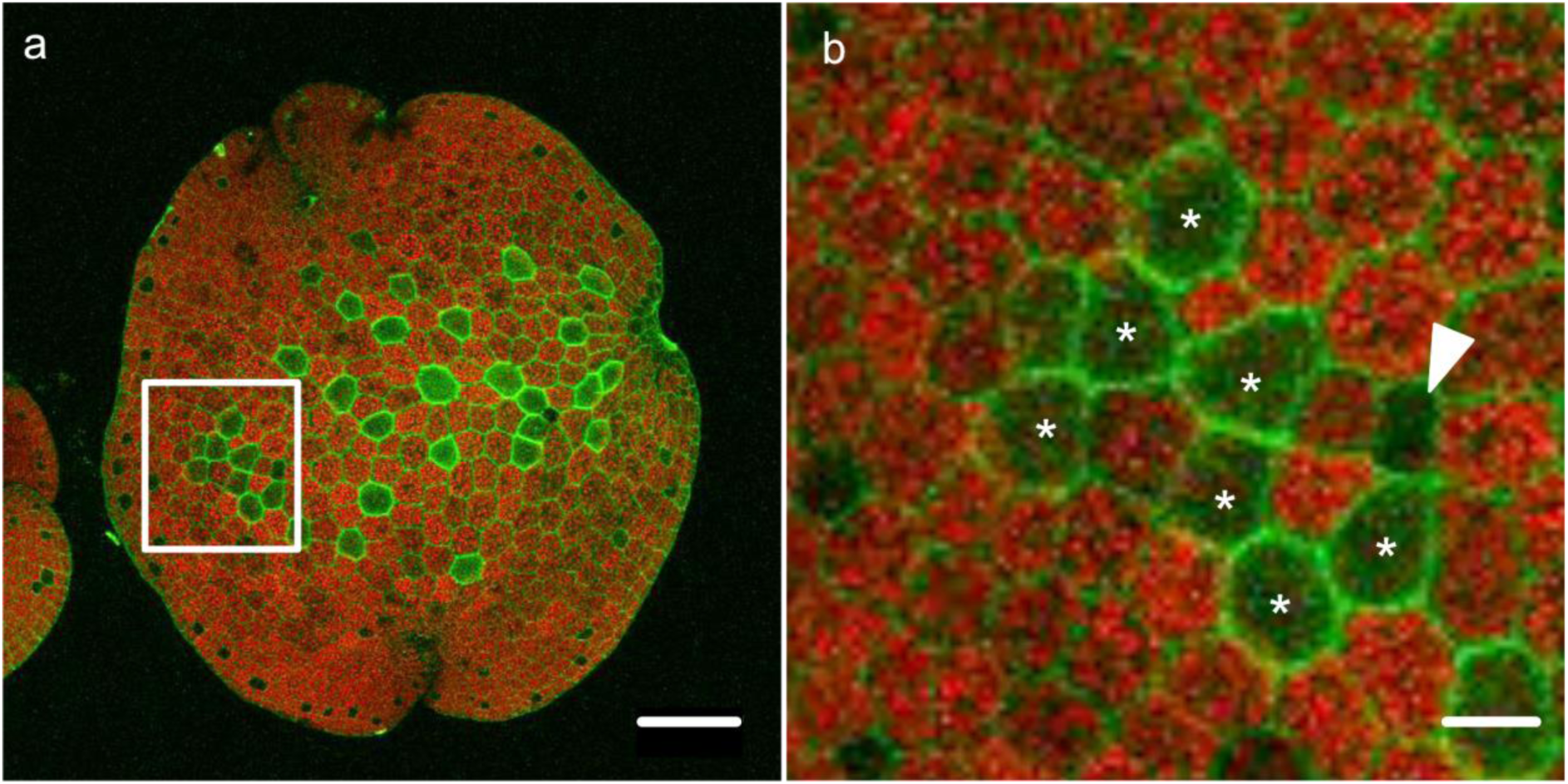
One 7-cell rhizoid cluster develop on Tak-2 gemma. a. Rhizoid patterning of a Tak-2 gemma showing individual rhizoid cells, smaller rhizoid cluster and a 7-cell rhizoid cell cluster (chlorophyll in red, PI in green, scalebar = 100 µm). b. Magnification of the 7-cell rhizoid cluster highlighted in a (scalebar = 20 µm).

**Supplementary Figure 2.**
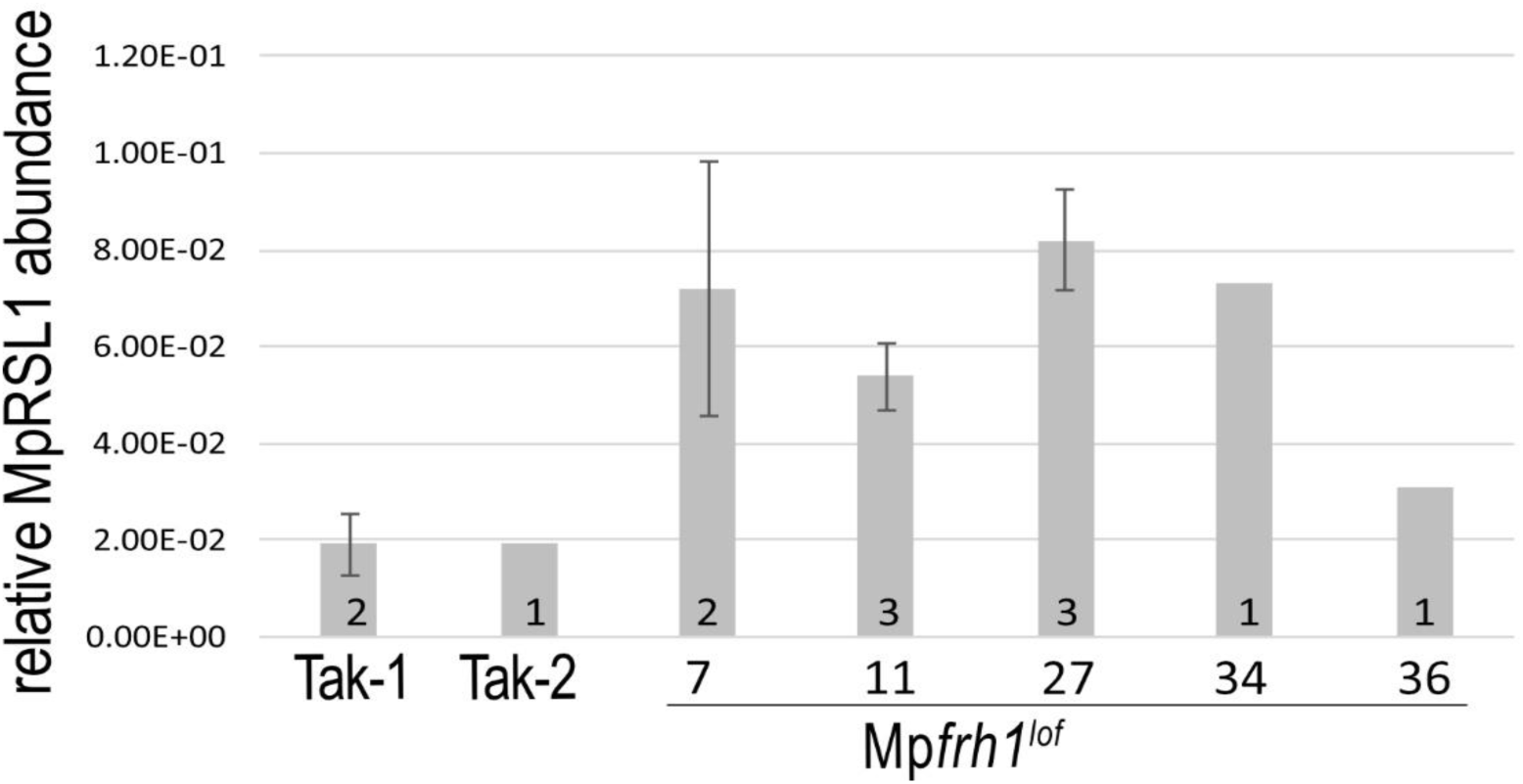
expression of MpRSL1 in Mpfrh1^lof^ mutants. Amplification of MpRSL1 mRNA from RNA isolated from in 7-day old thalli grown from gemmae in wild type (Tak-1, Tak-2) plants and Mp*frh1^lof^* mutants; mRNA levels were normalized to the housekeeping gene Mp*APT1*. n(Tak-1) = 2, n (Tak-2) = 1, n(Mp*frh1^lof-7^*) = 2, n(Mp*frh1^lof-11^*) = 3, n(Mp*frh1^lof-27^*) = 3, n(Mp*frh1^lof-34^*) = 1, n(Mp*frh1^lof-36^*) = 1.

**Supplementary Figure 3.**
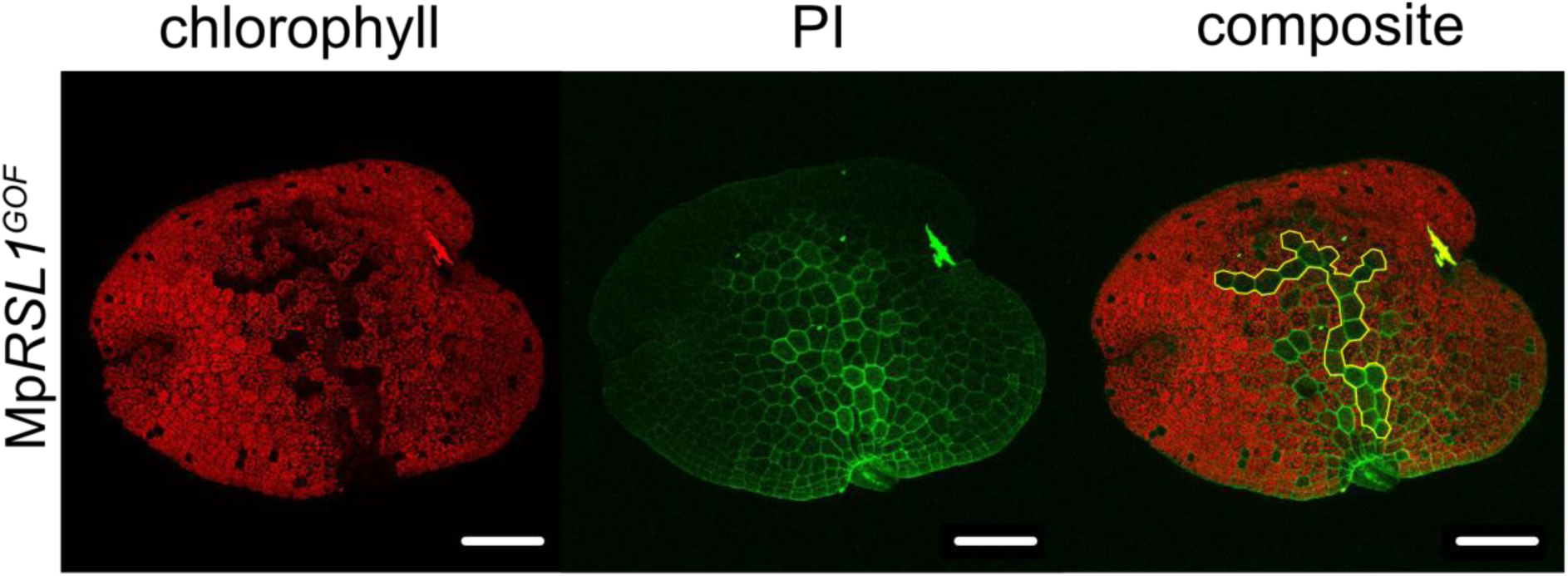
21-cell rhizoid cluster develop on Mp*RSL1^GOF^* gemma. Rhizoid patterning of a Mp*RSL1^GOF^* gemma showing a 21-cell rhizoid cell cluster highlighted as yellow perimeter (first panel = chlorophyll, second panel = PI, third panel = composite, scalebar = 100 µm).

